# Understanding the Transfer and Persistence of Antimicrobial Resistance in Aquaculture Using a Model Teleost Gut System

**DOI:** 10.1101/2024.07.30.605792

**Authors:** Alexandru Stefan Barcan, Joseph Humble, Sandeep Kasaragod, Mohammad Saiful Islam Sajib, Rares Andrei Barcan, Philip McGinnity, Timothy J. Welch, Brendan Robertson, Emanuel Vamanu, Antonella Bacigalupo, Martin Stephen Llewellyn, Francisca Samsing

## Abstract

The development, progression, and dissemination of antimicrobial resistance (AMR) is determined by interlinked human, animal, and environmental drivers, posing severe risks to human health. Conjugative plasmid transfer drives the rapid dissemination of AMR among bacteria. Besides antibiotic judicious use and implementation of antibiotic stewardship programs, mitigating antibiotic resistance spread requires an understanding of the dynamics of AMR transfer among microbial communities, as well as the role of various microbial taxa as potential reservoirs that promote long term AMR persistence. Here, we employed Hi-C, a high-throughput, culture-free technique, combined with qPCR, to monitor carriage and transfer of a multidrug-resistant plasmid within an Atlantic salmon in vitro gut model during florfenicol treatment, a benzenesulfonyl antibiotic widely deployed in fin-fish aquaculture. Microbial communities from the pyloric ceaca of three healthy adult farmed salmon were inoculated into three bioreactors developed for the SalmoSim gut system. The model system was then inoculated with an *Escherichia coli* strain ATCC 25922 carrying plasmid pM07-1 and treated with florfenicol at a concentration of 150 mg/L fish feed media for five days prior to a washout/recovery phase. Hi-C and metagenomic sequencing identified numerous transfer events, including to gram-negative and gram-positive taxa and, crucially, continuing transfer and persistence of the plasmid once florfenicol treatment had been withdrawn. Our findings highlight the role of commensal teleost gut flora as a reservoir for AMR, and our system provides a model to study how different treatment regimes and interventions may be deployed to mitigate AMR persistence.

## Main

Antimicrobial resistance (AMR) poses a critical challenge to global health. Unwinding the emergence, evolution, and transmission dynamics of individual antibiotic resistance genes (ARG) in microorganisms is essential in developing sustainable strategies against this threat (1). Facilitated by horizontal gene transfer (HGT), multidrug resistance (MDR) plasmids facilitate the rapid spread of ARGs in bacteria (2). This phenomenon is especially relevant in the microbiome of vertebrates’ guts, which accelerates the emergence of antibiotic-resistant bacterial strains (3). Aquaculture, one of the world’s fastest-growing food production industries, remains the least researched field in AMR (4). In this study, we used the Atlantic salmon artificial gut system, SalmoSim (Fig. S1), to track a MDR plasmid before, during and after treatment with an antibiotic widely deployed in finfish aquaculture.

Specifically, we aimed to establish: (i) the host range and adaptability of the MDR pM07-1 plasmid (5) within the complex microbial community of the Atlantic salmon gut, (ii) the impact of selective pressure exerted by the antibiotic florfenicol on the dynamics of plasmid transfer and the subsequent shifts in microbial community structure, and (iii) the role of different microbial taxa in promoting long term persistence of AMR determinants in the absence of drug pressure.

To address these questions, we collected the intestinal tracts from three healthy Atlantic salmon and extracted gut microbiome samples from their pyloric cecum segments. A custom fish feed medium was prepared (6, 7), and three 700 mL bioreactors were set up to simulate the salmon gut environment (Fig. S1). Each bioreactor was inoculated with a gut microbiome sample and maintained under anaerobic conditions. *Escherichia coli* ATCC 29522, harboring the multidrug-resistant plasmid pM07-1, was then inoculated into the system. The plasmid pM07-1 was originally isolated from the bacterium *Edwardsiella ictaluri*, found in a moribund *Ictalurus punctatus* suffering from enteric septicemia (5). The experimental trial included phases of static microbial growth, continous feed provission, plasmid inoculation (Pretreatment), antibiotic treatment (Treatment), and a depuration period (Washout) (Fig. S2). Bioreactor samples underwent qPCR with florfenicol resistance gene primers to quantify pM07-1 plasmid concentration, and Hi-C libraries were prepared with the Phase Genomics ProxiMeta Hi-C v4.0 Kit using the manufacturer-provided protocol (Fig. S3) (8). (For more Materials and Methods, see Supplementary Information).

Initial microbial diversity, represented by the ABC Mix sample from static growth (Fig. 1a) was high, as shown by the Shannon diversity index (Fig. 1b) and rarefaction curves (Fig. 1c). However, continuous flow with fish feed reduced microbial diversity prior to inoculation of the plasmid across all bioreactors. In Bioreactor A (Fig. 1a), there was a reduction in *Bacillus* and *Aliivibrio* abundance during the treatment and washout phases, with a concurrent increase in *Escherichia* and *Stenotrophomonas* populations, both carrying florfenicol resistance genes. Bioreactor B exhibited a rise in *Bacillus* and *Psychrobacter* abundances during the treatment and washout phase, with each group carrying the resistance factor suggesting that florfenicol selection enriched for bacterial genera harboring the *floR* resistance gene. In contrast, in Bioreactor C, the removal of the selective pressure during the washout phase led to a decline in the resistant *Psychrobacter* population and an increase in the susceptible *Bacillus, Photobacterium*, and *Aliivibrio* populations. The distinct clustering of samples by bioreactor in the Bray-Curtis NMDS plot (Fig. 1d) indicates that bioreactor, and thus individual host innoculaum, is a key determinant of microbial community composition. These differences could stem from the hosts’ genetic background, early-life environmental exposures, and unique host-microbe interactions.

**Fig. 1.**
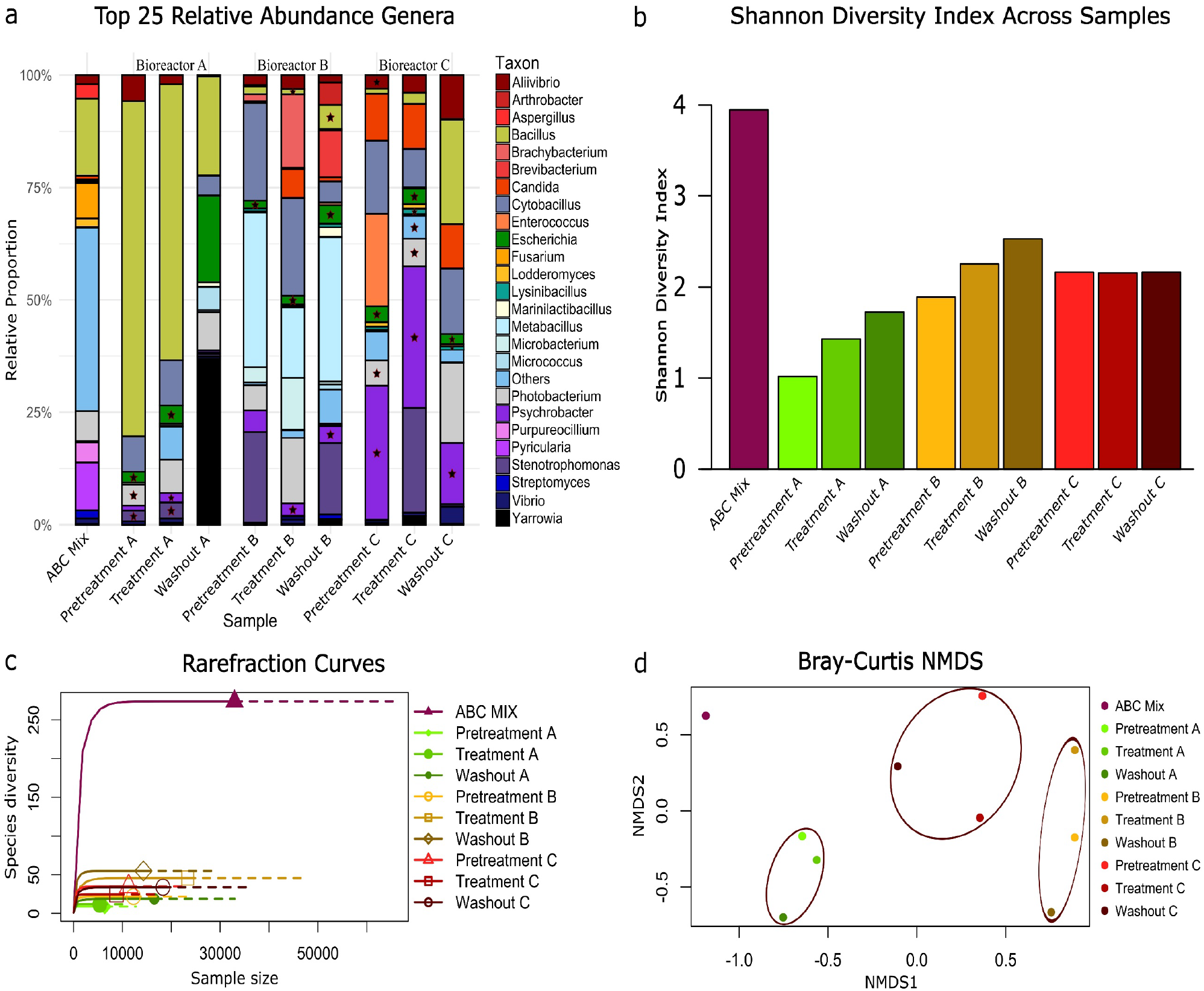
Species diversity and relative abundance of top 25 genera across all samples. **a**. Variation in relative abundance of top 25 taxa, across samples in different bioreactor phases. The ABC Mix bar, represents a composite from bioreactors A, B, and C after 5 days of static growth prior to the initiation of continuous flow with fish feed. Stars indicate genera with the florfenicol resistance gene. Diverse, less abundant genera are represented as “Others.” **b**. The Shannon Diversity Index. The Y-axis represents the Shannon Diversity Index, ranging from 0 to 4. **c**. The rarefaction curves show the ABC Mix sample which has the highest genera diversity, plateauing at 250 taxa. Other samples show lower diversity, below 50 species, with less pronounced plateaus, indicating lower richness. **d**. The Bray-Curtis NMDS plot highlights relationships between microbial community compositions across bioreactor samples. Brown ellipses encapsulate clusters, showing samples from the same bioreactor. The ABC Mix (dark purple) is distinct, indicating a unique microbial community.

During the washout phase of Bioreactor A, we observed a noticeable increase in *Yarrowia* populations (Fig. 1a) and none of the nine AMR genes associated with plasmid pM07-1 were detected (Fig. 2c). A mechanical failure of the magnetic stirrer responsible for maintaining bioreactor homogenization, which occurred five days before the last sampling point likely underpins this divergent profile. The mechanical failure resulted in an increase in pH to approximately 11. *Yarrowia*, a yeast commonly used as a probiotic in fish feed (9) and known for its high pH tolerance (10), thrived under these alkaline conditions. The fault which also appears to have purged many abundant bacteria, including those carrying the plasmid.

**Fig. 2.**
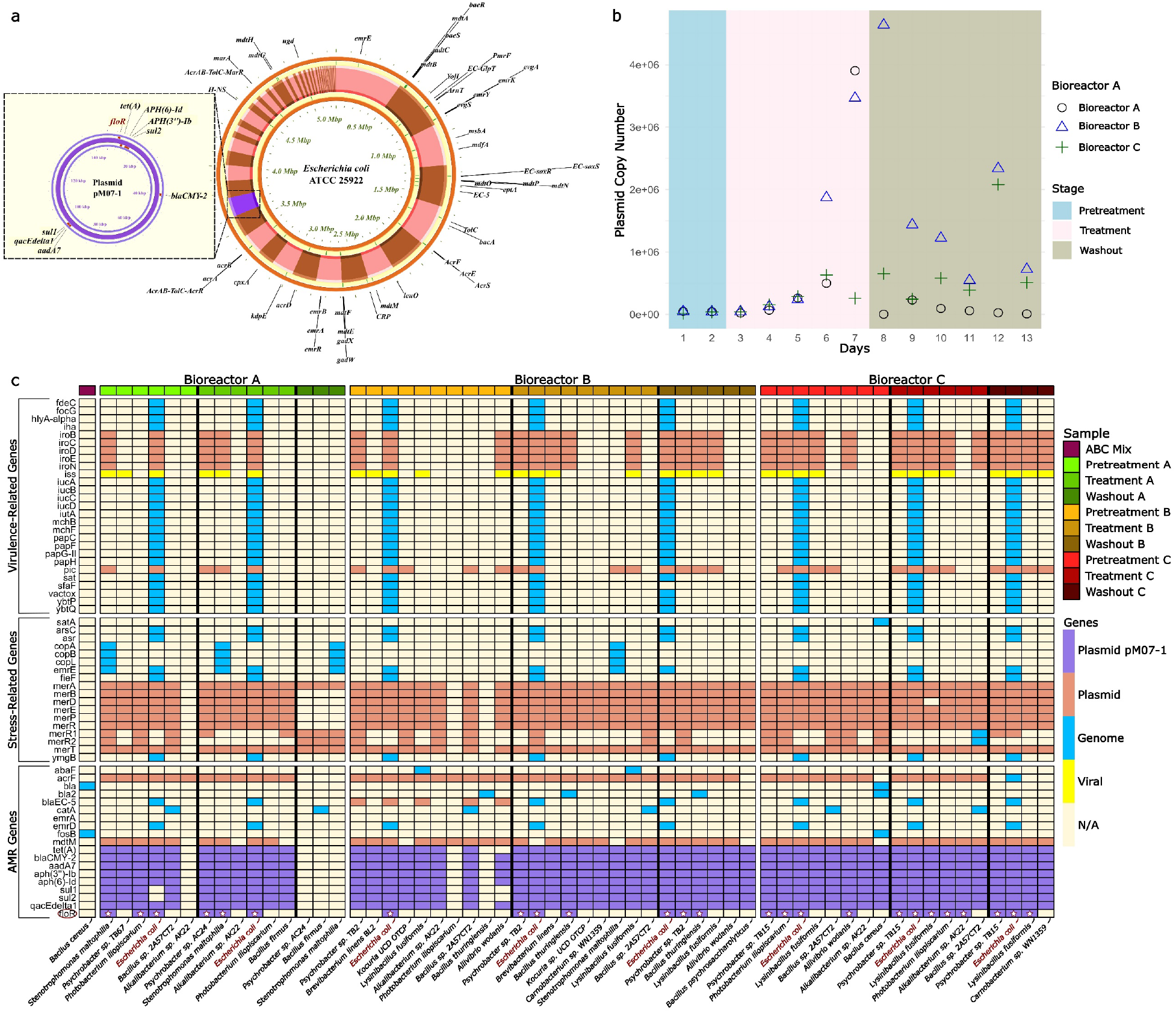
Plasmid distribution, concentration dynamics, and gene prevalence across all samples. a. Circular genome map of *Escherichia coli* strain ATCC 25922 with plasmid pM07-1. The map highlights antimicrobial resistance genes across the genome, emphasizing those on the approximately 150 kb plasmid pM07-1. Nine AMR genes are located on pM07-1, including the *floR* resistance gene highlighted in red. These genes are associated with resistance to various antibiotics: *floR* (chloramphenicol and florfenicol), *tet(A)* (tetracycline), *APH(6)-Id* and *APH(3’’)-Ib* (aminoglycosides), *sul2* and *sul1* (sulfonamides), *CMY-2* (carbapenems, cephalosporins, penicillins), *aadA7* (spectinomycin and streptomycin), and *qacEdelta1* (disinfecting agents and antiseptics). b. The concentration of plasmid copy numbers over 13 days in bioreactors A, B, and C was tracked during Pretreatment, Treatment, and Washout stages with qPCR. Plasmid copy numbers ranged from 0 to over 4 million copies/μl. Bioreactor A is shown with open circles, Bioreactor B with open triangles, and Bioreactor C with plus signs. During the pretreatment stage (Days 1-2), plasmid concentrations in all bioreactors were low and consistent. In the treatment stage (Days 3-7), bioreactor B saw a notable increase starting from Day 6, peaking at approximately 2 million copies/μl by Day 7. Bioreactor A also increased, peaking on Day 7 but lower than B. Bioreactor C had a less pronounced increase. During the washout stage (Days 8-13), bioreactor B’s plasmid concentrations continued to rise, peaking at 4 million copies/μl on Day 9 and fluctuating thereafter. Bioreactor A’s concentrations decreased to nearly zero by Day 13, while bioreactor C showed a modest increase, peaking around Day 11. c. Heatmap highlighting the presence of AMR genes, stress-related, and virulence-related across bioreactor samples A (green), B (brown), and C (red) during Pretreatment, Treatment, and Washout stages. Plasmid pM07-is shown in purple. *Escherichia coli*, the plasmid vector, consistently displayed all plasmid-specific AMR genes except in the ABC Mix and Washout A samples. Noticeable plasmid transfer occurred across various stages, with Bioreactor B showing the highest diversity and presence of AMR genes, particularly during Treatment and Washout.

Plasmid transfer from Gram-negative to Gram-positive bacteria is rare, and when reported typically involves shuttle, a type of plasmid that has the ability to replicate in multiple host species (11, 12) or broad host range plasmids like IncP (13) and IncPromA (14), with narrower host plasmids like IncX3 also showing cross-Gram transfer (15). While previous studies have reported inter-Gram plasmid transfer for other plasmid groups, our findings demonstrate that the MDR IncA/C plasmid pM07-1 can also traverse the Gram-negative/Gram-positive boundary.

We identified plasmid transfer to 16 different bacterial species, comprising 7 Gram-negative and 9 Gram-positive species (Fig. 2c). These species belong to seven different bacterial families, demonstrating the broad host range and adaptability of the plasmid. The Gram-negative species included members of the families *Vibrionaceae, Moraxellaceae*, and *Xanthomonadaceae*, while the Gram-positive species were distributed among the families *Bacillaceae, Carnobacteriaceae, Brevibacteriaceae*, and *Micrococcaceae*.

Sequencing of the first Hi-C sample – the combined innoculum (ABC mix), identified only two AMR genes, *bla*, and *fosB*, on *Bacillus cereus* chromosomes (Fig. 2c). These genes are associated with resistance to beta-lactams and Fosfomycin (16), and indicate that some AMR genes are in circulation in farmed Atlantic Salmon, but not apparently those that might correspond to the frontline antimicrobials currently deployed in a veterinary context (i.e. tetracycline, florfenicol) (17).

Hi-C is a powerful metagenomic tool but has limitations that could affect data interpretation. Inconsistencies in detecting AMR genes like *floR, sul1*, and *sul2* across species (Fig. 2c) can result from insufficient sequencing depth (18), challenges in mapping short reads (19), and variability in region capture (20), impacting gene detection consistency across samples. Fig. 2c shows noticeable HGT of pM07-1 across diverse bacterial species, even during the pretreatment stage when plasmid concentrations were uniformly low (Fig. 2b). This early HGT activity, demonstrated by the presence of AMR genes in multiple species, demonstrates the persistent nature of HGT in microbial ecosystems and suggests that minimal plasmid presence can facilitate the spread of resistance genes, even in the absence of strong selective pressure.

Antibiotic selective pressure during the last 2 days of treatment phase significantly increased plasmid abundance across all bioreactors. Specifically, plasmid abundance increased significantly in all observations during the treatment phase compared to the pretreatment phase (p < 0.05). Notably, this increase corresponded with the highest diversity of AMR gene presence and the introduction of new bacterial species harboring these genes, such as *Brevibacterium linens* and *Kocuria UCD OTCP* (Fig. 2c). This indicates that antibiotic exposure not only increased plasmid concentration but also promoted active plasmid transfer and establishment in novel bacterial hosts.

Furthermore, there was no significant decrease in plasmid abundance during the washout phase, with 2 out of 3 observations showing no significant change compared to the treatment phase (p > 0.05). This suggests a persistence of increased plasmid levels post-treatment, indicating that antibiotic exposure not only elevates plasmid concentration but also potentially facilitates the retention of these plasmids even after the removal of antibiotic pressure (Fig. 2b).

In summary, our findings revealed that: (i) plasmid pM07-1 has a broad host range, transferring to 16 bacterial species, of both Gram-negative and Gram-positive bacteria, (ii) under florfenicol selective pressure, plasmid transfer increased, leading to higher plasmid concentrations and shifts in microbial community structure, favoring resistant strains while reducing diversity among susceptible populations. (iii) plasmid transfer and maintenance seem to have little cost, can occur in the absence of selection pressure, and persists in a wide variety of microbial taxa in the absence of selection.

## Acknowledgements

The authors would like to acknowledge the dedicated staff at the OHRBID lab for their invaluable support and contributions: Taya Forde, Ryan Carter, Jo Halliday, Katarina Oravcova, and Mary. Additionally, special thanks to Raminta Kazlauskaite and Lajos Kalmar for their for their assistance.

## Ethics declarations

This experiment was conducted at the biological safety category 2 OHRBID laboratory and used SOPs and Risk Assessments which were approved by the University of Glasgow’s biological safety committee.

## Data Availability Statement

The data supporting the findings of this study have been deposited in the NCBI Sequence Read Archive (SRA) under BioProject accession number PRJNA1135464.

