## Supplementary Materials and Methods for "Understanding the Transfer and Persistence of Antimicrobial Resistance in Aquaculture Using a Model Teleost Gut System"

### Atlantic salmon gut samples collection and preparation

Three intestinal tracts were collected from healthy farmed adult Atlantic salmon. They were transported to the laboratory in an anaerobic box on ice to maintain their integrity and prevent microbial contamination. Upon arrival at the laboratory, the pyloric cecal segments were carefully separated from each intestinal sample. A total of 1 g of gut microbial sample (comprising mucus and scraping from the internal epithelium) was extracted from each pyloric cecum and transferred into separate sterile cryovial tubes. Special care was taken to avoid cross-contamination between the individuals. The gut microbial samples were snap-frozen in liquid nitrogen and stored in a freezer at a temperature of -70°C to seed the bioreactors as described below. In addition, we collected a set of "backup" samples to serve as a contingency plan to maintain the experimental integrity and allow for potential restarts, if required.

### Fish feed medium preparation

The feed pellets used for the primary in vitro culture system had the nutritional information as previously described (1, 2). To prepare the feed medium, a mixture of 70 g of Instant Ocean and 20 g of ground fish pellets were combined with 2 L of Milli-Q water in a Duran bottle. To simulate the enzyme digestion of salmon's stomach, the pH of the solution was adjusted to 3.5 using hydrochloric acid (HCl) and 12 ml of crude stomach enzyme extracts were added while stirring. The enzyme digestion process was carried out for one hour. Subsequently, the pH was restored to neutral by adding sodium hydroxide (NaOH). Once the pH reached 7.0, 1 g of dry sterile mucus was added to the digested media, and the entire mixture was autoclaved at 121 °C for 15 min to ensure sterility. After autoclaving, the fish meal was filtered using a sieve to remove any solid particles that could block the silicon tubes and then autoclaved once again. Each bioreactor was provided with its batch of fish feed medium for the experiment.

### SalmoSim in vitro system preparation.

Three 700-mL custom-made double-jacketed glass bioreactors was used in this study. To support bacterial growth four pieces of 1 $cm^3$ aquarium sponge filter werewas placed in each bioreactor as a substrate. Magnetic stirrers were used for continuous mixing. The entire setup, including the bioreactors and the magnetic beads, was autoclaved to ensure sterility. A microcontroller board (Arduino Uno) was used to monitor temperature and control and monitor pH via automated pumping of acid (0.1 M hydrochloric acidHCl) and base (0.1 M NaOH) buffers (see Fig. S1). pH buffers and feed media were pumped using Atlas Scientific peristaltic pumps (Atlas Scientific) controlled by the microcontroller board.  Arduino. Initially, 400 mL of sterile feed media was added to each bioreactor. During the continuous flow phases of the experiment, fish feed was supplied at a rate of 200 ml/day, with waste slurry being removed by a peristaltic pumping to keep a constant total volume of 400 ml/bioreactor.

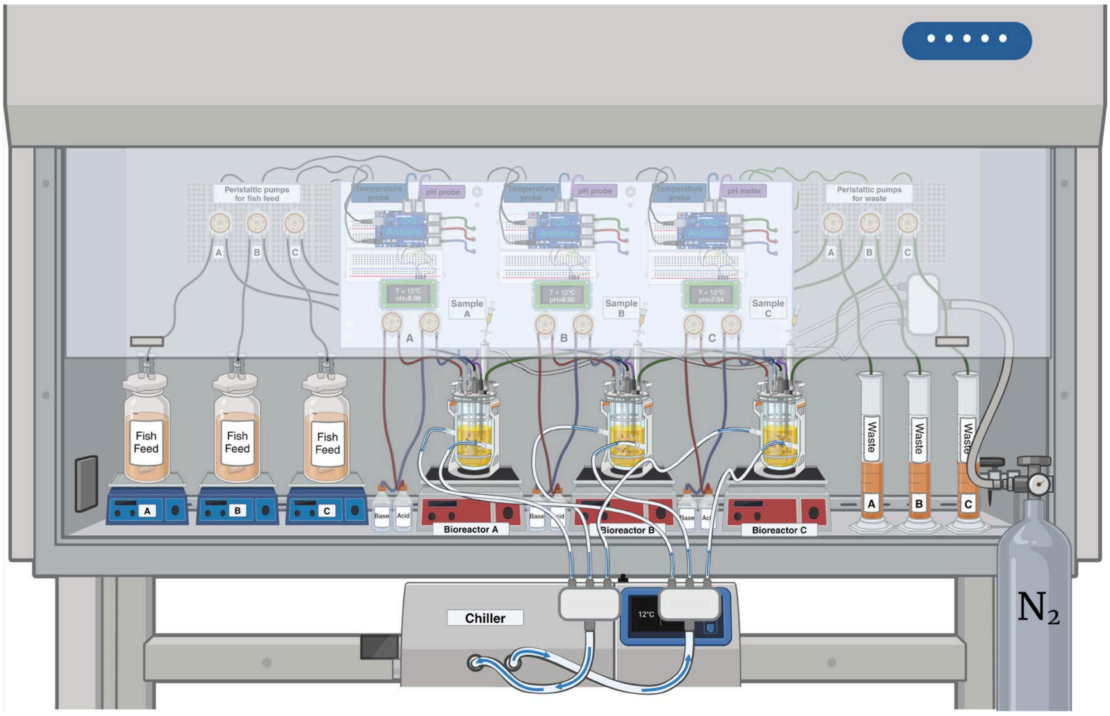


**Fig. S1 Experimental Setup**

Fig. S1 Illustrates the SalmoSim in vitro system configured within a biosafety cabinet. The system includes three 700-mL custom-made double-jacketed glass bioreactors labelled A, B, and C, each equipped with magnetic stirrers for continuous mixing and containing four pieces of 1 $cm^3$ aquarium sponge filters to support bacterial growth. Temperature and pH are meticulously controlled by a microcontroller board (Arduino Uno), which regulates automated dosing of acid and base buffers (0.1 M hydrochloric acid and 0.1 M NaOH) buffers using Atlas Scientific pumps to regulate pH. Initial sterile feed media of 400 mL was introduced into each bioreactor. Feed media was continuously supplied at 200 mL/day through peristaltic pumps, while waste was simultaneously removed to maintain a stable volume of 400 mL per bioreactor. A chiller was integrated to sustain the system at 12°C, and a nitrogen ($N_2$) tank ensures anaerobic conditions were maintained. The detailed wiring and tubing setup depicted in the figure ensured precise monitoring and control, crucial for studying antimicrobial resistance in microbial communities within the artificial gut environment of Atlantic salmon. This setup facilitates a comprehensive investigation of bacterial dynamics under controlled laboratory conditions.

### Inoculation and Maintenance of Bioreactors:

Each bioreactor was inoculated with 1 g of gut contents containing intestinal microorganisms from the gut microbiome sample from the pyloric cecum, dissolved in 1 mL of autoclaved 35 g/L Instant Ocean Sea Salt suspension. This inoculum e gut mucus served as the source of intestinal microorganisms, representing the microbial composition of the fish Atlantic salmon’s pyloeric caecum . This step ensured the establishment of a representative microbial community in each bioreactor. The internal physiological and chemical environments of the bioreactors were carefully controlled to mimic in vivo conditions. The dissolved oxygen level in the system was intended to be minimized by sparging the bioreactors with sterile-filtered nitrogen (N2) gas for 20 minutes daily. The temperature of the bioreactors was regulated at 12 °C by pumping chilled water through the outer compartment of the double-jacketed bioreactors. To simulate the pyloric cecum compartments the three bioreactors received daily additions of 1 mL of filtered salmon bile and 0.5 mL of autoclaved 5% mucous solution. The 5% mucous solution was prepared by mixing 5 g of mucous with 100 mL of Milli-Q water, which was then autoclaved to maintain sterility and aliquoted in 1.5 ml Eppendorf tubes and frozen at -20 °C. This experimental setup ensured the provision of appropriate nutrition and environmental conditions, mimicking the pyloric cecum compartments of the fish. The continuous monitoring and control of pH, temperature, and nutrient additions allowed for the maintenance of a stable and representative in vitro fish gut system.

**Bacterial Strains and Plasmid Transfer**

The strain used in this study was *Escherichia coli* ATCC 29522, deficient of diaminopimelic acid (DAP) and harboring a multidrug-resistant plasmid pM07-1 (3, 4). In order to avoid the continuous use of DAP, the plasmid was transferred to *Escherichia coli* ATCC 25922 (5), via conjugation, which was performed according to the general conjugation protocol from Barrick Lab (6). This transfer step was carried out prior to the preparation of the inoculum for the subsequent experiments.

### Plasmid Innoculum Preparation

Florfenicol stock solution was prepared by dissolving florfenicol in dimethyl sulfoxide (DMSO) at a concentration of 50 mg/mL. The stock solution was filter sterilized in a 0.2 µm filter syringe and stored as aliquots at -20 °C.

Before inoculating the gut simulator, ATCC 25922 harboring pMO7-1 needed to be acclimatized to the SalmoSim system conditions. Initially, it was cultured overnight in tryptic soy broth supplemented with florfenicol at a concentration of 50 µg/mL. The cultures were incubated at 37°C with agitation at 300 rpm. To simulate the conditions of the bioreactors, a customized marine media supplemented with florfenicol similar to the media used in the bioreactors was prepared. The ATCC 25922(pMO7-1) strain was transferred to this customized media, which also contained florfenicol. The culture was grown overnight at 25°C with agitation at 300 rpm. Subsequently, ATCC 25922(pMO7-1) was cultured in fish feed media supplemented with florfenicol (50 mg/L), and the culture was incubated at 14 °C for 40 hours.

To prepare the inoculum, ATCC 25922(pM07-1) grown in the fish feed media was centrifuged at 2000 g for 10 minutes. The resulting pellet was washed with sterile seawater to remove the antibiotic. After washing, the pellet was suspended in 1 mL of seawater. The system was then inoculated with approximately 10^10 colony-forming units (CFU).

### Experimental Design and Sampling

The experimental trial followed a predetermined timeline involving static microbial growth, fish feed supplementation, plasmid inoculation, antibiotic treatment, and a subsequent washout period. Sampling was conducted at specific time points throughout the study for analyses such as Hi-C, and qPCR. Initially, the experiment involved five days of static microbial growth without the continuous flow of fish feed media, allowing bacterial populations to establish within the in vitro system. This was followed by a 35-day period of fish feed supplementation at 200 mL/day, supporting bacterial proliferation and activity. On day 40, the bioreactors were inoculated with plasmid-conjugated *E. coli*. Media supplementation was paused for 24 hours post-inoculation to allow the conjugated bacteria to settle, after which continuous flow of feed media and waste removal resumed. This step introduced transmissible genetic material into the bacterial populations to study the spread of antimicrobial resistance.

Starting on day 42 florfenicol was added to the fish feed media at 150 mg/L for five days, providing a selective pressure to stimulate plasmid transfer between bacterial communities. This was followed by a seven-day washout period without antibiotic treatment.

Samples were collected at various stages to capture different phases of the experiment. Hi-C sequencing samples were taken on day 5, and additional samples were collected on day 42 before adding florfenicol (Pretreatment), on day 47 (the last day of antibiotic Treatment phase), and on day 54 (end of study, post-washout phase). Additionally, samples for qPCR analysis were collected daily, starting from day 39 (Fig. S2).

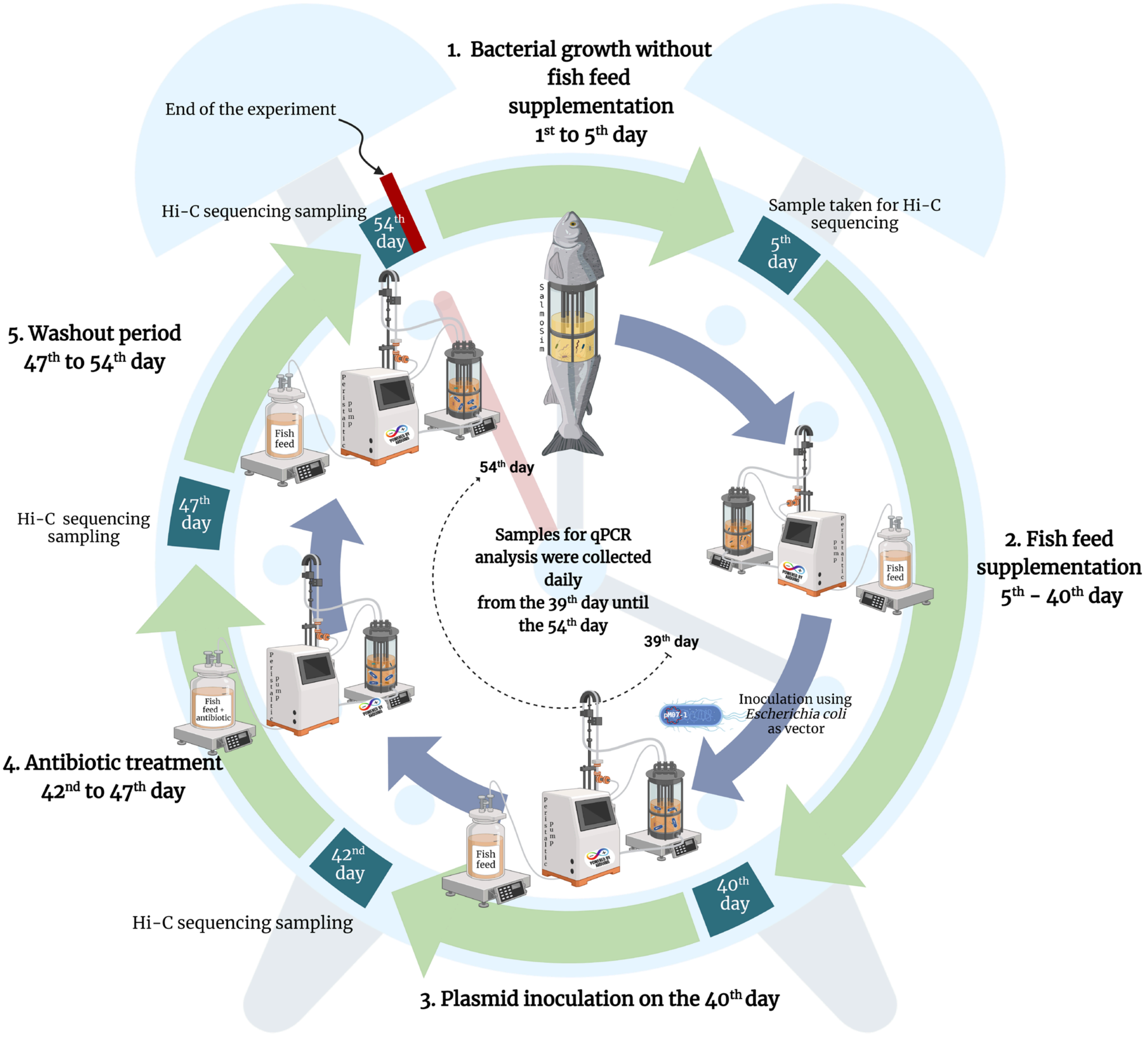


**Fig. S2 Experimental Design and Sampling**

### Analyzing the pM07-1 plasmid concentration during the main experimental trial

During the experimental trial, 1 mL of each bioreactor sample was collected in sterile 1.5 mL Eppendorf tubes. The collected samples were then subjected to centrifugation at 1000 g for 10 minutes at 15 °C. Subsequently, the supernatant was carefully removed, and the resulting pellet was suspended in 1 mL of sterile water. To facilitate plasmid extraction the samples were boiled for 15 minutes. After boiling, the samples were centrifuged at 13,000 g for 10 minutes to separate the supernatant. To quantify plasmid levels in each bioreactor, we used a The quantitative real time PCR (qPCR) analysis was performed using 0.5 mL of the recovered supernatant obtained after centrifugation. Primers targeting the florfenicol resistance gene were used to determine levels of the multidrug-resistant plasmid pM07-1: The specific target gene associated with plasmid pM07-1, the presence of the florfenicol resistance gene was quantified using qPCR with the following primers: flor-q-FP GGATGGCAGGCGATATTCAT and flor-q-RP CTTGACTTGATCCAGAGGGC. The pM07-1 plasmid was amplified and purified using the QIAGEN Plasmid Plus Midi Kit in order to create standards of known concentration. This was done so that a standard curve could be generated to enable absolute quantification of experimental samples by qPCR. Serial dilutions were prepared from the purified plasmid product as follows: P1 (1 in 100), P2 (1 in 1000), P3 (1 in 10,000), P4 (1 in 100,000), P5 (1 in 1,000,000), and P6 (1 in 10,000,000). The qPCR reaction was performed using Luna® Universal qPCR Master Mix (BioLabs®, New England) at a concentration of x 1x. The florfenicol resistance qPCR primers were used at a working concentration of 0.25 μM. 2 μL of either the experimental sample or plasmid standard were used in each reaction and the final total volume was 10 μL. Each sample was analyzed in technical duplicates. The qPCR analysis was performed in Qiagen Strip Tubes using the a Qiagen qPCR machine (Rotogene Q).

The plasmid copy number in each sample was calculated by interpolating the Ct values from the bioreactor samples onto the standard curve equation. The copy number was expressed as copies/μl of the sample. The standard curve equation used for this calculation was derived from a linear regression of the known plasmid concentrations and their corresponding Ct values, using the following equation:

$$\text{Log (Copy Number)} = a\times(Ct)+b,$$

where a and b are the slope and intercept of the standard curve, respectively. This equation allows for the determination of plasmid copy numbers in the experimental samples by substituting the Ct values into the equation and solving for the copy number.

### Sampling preparation for Hi-C sequencing

10 mL from each bioreactor were collected on days 5, 42, 47, and 56 in 15 mL sterile tubes, then centrifuged at 3000 g for 10 min. The supernatant was discarded, and the pellet was resuspended in 0.5 mL of Phase Genomics PGShield™. Subsequently, the samples were sent to Phase Genomics (Seattle, WA, U.S.A.) for further analysis (see Fig. S3).

### ProxiMeta Methods

A Hi-C library was created with the Phase Genomics ProxiMeta Hi-C v4.0 Kit using the manufacturer-provided protocol (7). Briefly, intact cells were crosslinked using a formaldehyde solution, simultaneously digested using the Sau3AI and MlucI restriction enzymes, and proximity ligated with biotinylated nucleotides to create chimeric molecules composed of fragments from different regions of genomes that were physically proximal in vivo. Proximity-ligated DNA molecules were pulled down with streptavidin beads and processed into an Illumina-compatible sequencing library. Separately, using an aliquot of the original sample, DNA was extracted with a ZYMObiomics DNA miniprep kit (8), and a metagenomic shotgun library was prepared using ProxiMeta library preparation reagents. Sequencing was performed on an Illumina NovaSeq generating PE150 read pairs for both Hi-C and shotgun libraries. Hi-C and shotgun metagenomic sequencing files were uploaded to the Phase Genomics cloud-based bioinformatics portal for subsequent analysis (see Fig. S3A).

Shotgun reads were filtered and trimmed for quality and normalized using fastp (9) and then assembled with MEGAHIT (10, 11) using default options (Fig. S3B). Hi-C reads were then aligned to the assembly following the Hi-C kit manufacturer's recommendations (12). Briefly, reads were aligned using BWA-MEM (13) with the -5SP options specified, and all other options default. SAMBLASTER (14) was used to flag PCR duplicates, which were later excluded from the analysis. Alignments were then filtered with samtools (15) using the -F 2304 filtering flag to remove non-primary and secondary alignments. Metagenome deconvolution was performed with ProxiMeta (16, 17), resulting in the creation of putative genome and genome fragment clusters. Clusters were assessed for quality using CheckM (18) and assigned preliminary taxonomic classifications with Mash (19).

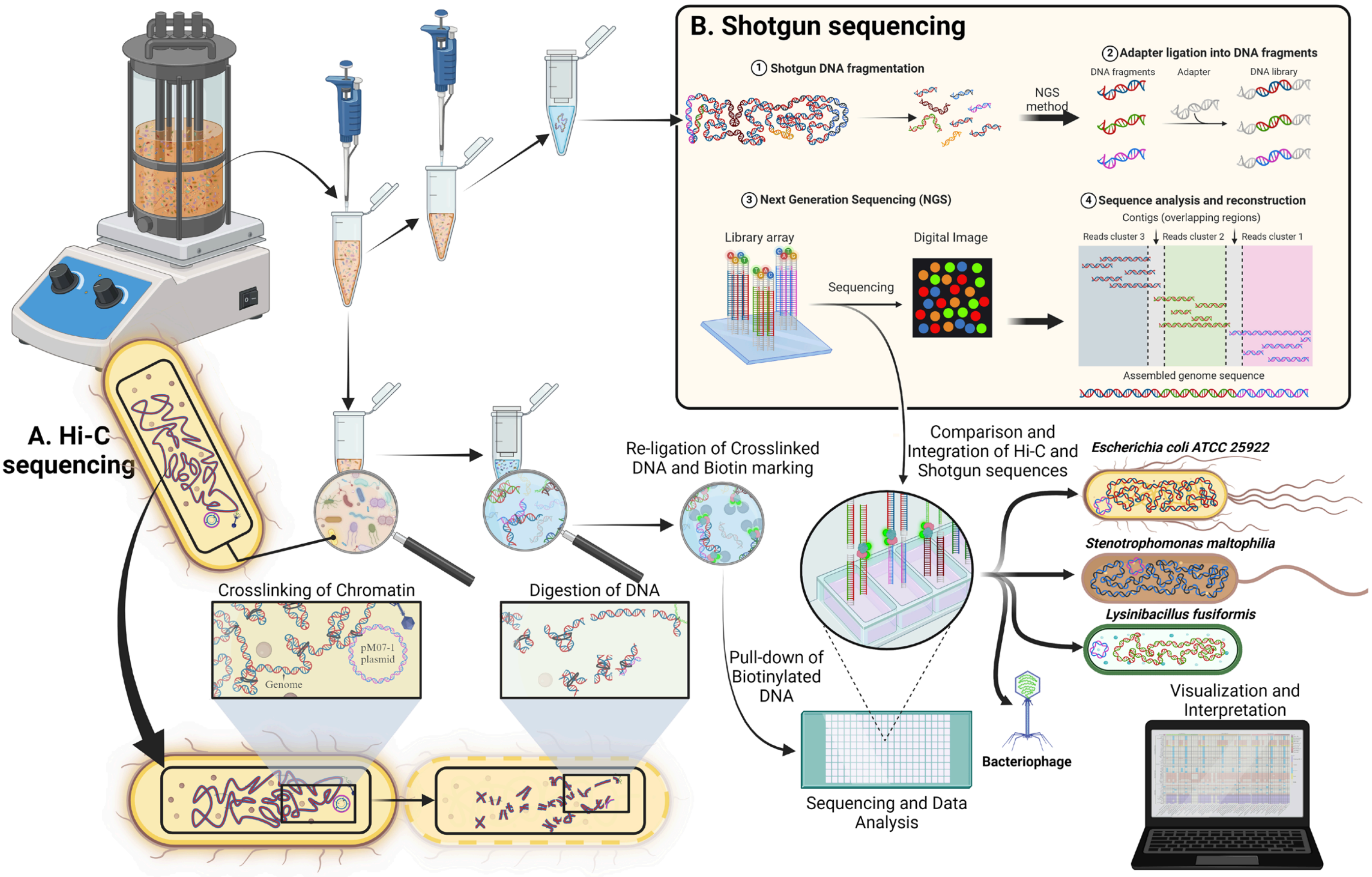


**Fig. S3 Integrated Hi-C and Shotgun Sequencing Analysis of Microbial Samples from Bioreactors**

In Fig. S3A the Hi-C sequencing process involves crosslinking chromatin within bacterial cells, followed by digestion of the DNA and re-ligation of the crosslinked fragments, which are then biotin-labelled. These labelled fragments are pulled down and sequenced, providing spatial information about DNA interactions within the cell. Simultaneously, in Fig. S3B, shotgun sequencing involves fragmenting DNA, ligating adapters, and using next-generation sequencing (NGS) to generate a library of sequences. This method allows for the comprehensive analysis and reconstruction of the bacterial genome. The integration of Hi-C and shotgun sequencing data enables a detailed comparison and comprehensive understanding of the microbial community's genetic structure and interactions. The final visualization and interpretation of the data reveal insights into the genomic organization and potential antimicrobial resistance mechanisms in bacterial species such as *Escherichia coli*, *Stenotrophomonas maltophilia*, and *Lysinibacillus fusiformis*.

### Taxonomic profiling

For the contigs generated from Phase Genomics, taxonomic profiling of the microbial communities present was conducted using Kraken version 2.1.2, which leverages a specialized microbial database to enhance the accuracy and specificity of the identification process. Subsequent estimation of species abundance was performed with Bracken, applying a threshold parameter of 100 (-r) and configured to exclusively identify taxa at the genus level.

Following the taxonomic profiling, microbial abundance data for each sample was visualized in RStudio. This involved the use of packages such as tidyr, dplyr, and ggplot2. To facilitate the calculation of abundances, data was transformed to long format. The top 25 taxa were identified by summing their abundances, and these taxa, along with an aggregated "Others" category, were plotted to highlight their relative proportions.

#### Whole Genome Sequencing, Contig Assembly, and Annotation

The *Escherichia coli* strain ATCC 25922 was isolated from the bioreactor inoculum, and its DNA was extracted using the Qiagen REPLI-g Single Cell Kit, following the manufacturer's instructions. The extracted DNA was sent to the company GEEWIZ for whole-genome sequencing. Genome assembly was conducted using SPAdes (20). Annotation of antimicrobial resistance genes was performed using the Comprehensive Antimicrobial Resistance Database (CARD) (21) through the PROKSEE server (https://beta.proksee.ca/projects). The PROKSEE server also generated high-quality, navigable maps for the whole genome's AMR genes, as well as for the plasmid PM07-1 and its AMR genes.
